# Genome variation and conserved regulation identify genomic regions responsible for strain specific phenotypes in rat

**DOI:** 10.1101/169748

**Authors:** David Martín-Gálvez, Denis Dunoyer de Segonzac, Man Chun John Ma, Anne E Kwitek, David Thybert, Paul Flicek

**Affiliations:** European Molecular Biology Laboratory, European Bioinformatics Institute, Wellcome Genome Campus, Hinxton, Cambridge, CB10 1SD, United Kingdom; Department of Pharmacology, University of Iowa, Iowa City, IA, USA; Iowa Institute of Human Genetics, University of Iowa, Iowa City, IA, USA

**Keywords:** Metabolic syndrome, genome regulation, evolution

## Abstract

The genomes of laboratory rat strains are characterised by a mosaic haplotype structure caused by their unique breeding history. These mosaic haplotypes have been recently mapped by extensive sequencing of key strains. Comparison of genomic variation between two closely related rat strains with different phenotypes has been proposed as an effective strategy for the discovery of candidate strain-specific regions involved in phenotypic differences.

We developed a method to prioritise strain-specific haplotypes by integrating genomic variation and genomic regulatory data predicted to be involved in specific phenotypes. To identify genomic regions associated with metabolic syndrome, a disorder of energy utilization and storage affecting several organ systems, we compared two Lyon rat strains, LH/Mav which is susceptible to MetS, and LL/Mav, which is susceptible to obesity as an intermediate MetS phenotype, with a third strain (LN/Mav) that is resistant to both MetS and obesity. Applying a novel metric, we ranked the identified strain-specific haplotypes using evolutionary conservation of the occupancy three liver-specific transcription factors (HNF4A, CEBPA, and FOXA1) in five rodents including rat.

Consideration of regulatory information effectively identified regions with liver-associated genes and rat orthologues of human GWAS variants related to obesity and metabolic traits. We attempted to find possible causative variants and compared them with the candidate genes proposed by previous studies. In strain-specific regions with conserved regulation, we found a significant enrichment for published evidence to obesity—one of the metabolic symptoms shown by the Lyon strains—amongst the genes assigned to promoters with strain-specific variation.

Our results show that the use of functional regulatory conservation is a potentially effective approach to select strain-specific genomic regions associated with phenotypic differences among Lyon rats and could be extended to other systems.

## Introduction

Phenotypic diversity is ultimately driven by genetic differences. The connections between DNA sequence and observed phenotypes are often difficult to determine and may be confounded by non-genetic causes including environmental effects. Regardless, it is increasingly clear that differences in transcriptional regulation are an important factor explaining phenotypic diversity (Wittkopp and Kalay 2012; Pai and Gilad 2014; Villar et al. 2014; Villar et al. 2015; Lowdon et al. 2016; Mack and Nachman 2016). This is especially true between closely-related species (Romero et al. 2012; Shibata et al. 2012; Stefflova et al. 2013; Pai and Gilad 2014). Accordingly, a number of efforts have been made to combine transcriptional regulatory data with genome variation to select candidate genomic regions involved in producing phenotypic characteristics of interest (Ward and Kellis 2012; Nica and Dermitzakis 2013; Lowe and Reddy 2015; Moreno-Moral and Petretto 2016).

The rat is a key animal model for biomedical research (Lindsey and Baker 2006; Jacob 2010; Aitman et al. 2016). More than 600 laboratory rat strains have been created over the last century in order to study specific traits including those which are more informative in rat than in other model species, such as behaviour and neurodegenerative diseases, cardiovascular diseases and metabolic disorders (Mashimo and Serikawa 2009; Voigt 2010; Yau and Holmdahl 2016). One focus over the last decade has been the identification of genes and other genomic loci associated with these strain-specific traits (Dwinell et al. 2011; Moreno-Moral and Petretto 2016). Despite the great number of quantitative trait loci (QTL) identified in rat models using a number of techniques (Shimoyama et al. 2014), only a small number of causative genes have been determined for complex traits or diseases (Aitman et al. 2010; Baud et al. 2013; Moreno-Moral and Petretto 2016).

Most genomic variants in an individual are expected to be neutral, and therefore have no impact on reproduction or survival (Kimura 1968; King and Jukes 1969). In the case of laboratory rats, the existing variation among strains (e.g. Hermsen et al. 2015) is the sum of the ancestral variation among individuals used in the process of strain development and the novel variation that originated and accumulated in the genome during the establishment and maintenance of the strains. Like humans (1000 Genomes Project Consortium, 2015) and laboratory mice (Adams et al. 2015), genetic variation among rat strains is not randomly distributed across the genome; instead it is organised in haplotype blocks (Saar et al. 2008; Atanur et al. 2013; Ma et al. 2014), which are caused by meiotic crossover of the shared ancestral variation. Comparison of these haplotype blocks among rat strains with different phenotypes has proven to be a powerful strategy for genetic mapping of complex traits and diseases (Saar et al. 2008; Ma et al. 2014). For example, Atanur and colleagues analysed the genomes of 27 rat strains, and found that haplotype blocks with variants that are unique to a single strain were positively selected in the initial phenotype-driven derivation of these strains, and thus variants associated with strain-specific phenotypes are predicted to be in these regions (Atanur et al. 2013). However, the genomic extent of such regions and the number of sequence variants found within them are nearly always too large for an effective determination of candidate loci influencing the phenotype of interest (see e.g. Cuppen 2005).

Regulatory activity such as active promoters, enhancers and transcription factor binding sites (TFBS) can be effectively mapped genome-wide with current techniques such as chromatin immunoprecipitation followed by high-throughput sequencing (ChIP-seq) (Encode Project Consortium et al. 2012). Previous studies have suggested that both the number and conservation level of transcription factor binding sites in a given region affect the level of gene expression (Pennacchio and Rubin 2001; Berman et al. 2002; Cheng et al. 2014; Villar et al. 2014; Wong et al. 2015). Since tissue characteristics are directed to a large extent by the activity of tissue-specific transcription factors, the location of these regulatory elements might be useful when selecting haplotype blocks associated with specific phenotypes or diseases.

In this study, we characterise the haplotype blocks holding strain-specific genome variation among three closely related strains of the Lyon rat. Although Lyon rats were initially established as a model of hypertension (Dupont et al. 1973), several additional symptoms related to metabolic syndrome (MetS), such as obesity, dyslipidaemia and susceptibility to insulin resistance have been found in the Lyon Hypertensive (LH/Mav) strain (Sassolas et al. 1981; Vincent et al. 1993; Wang et al. 2015). Only obesity is observed in the Lyon Low pressure (LL/Mav) and all MetS related phenotypes are absent in the Lyon Normotensive (LN/Mav) strain (Sassolas et al. 1981; Vincent et al. 1993; Bilusic et al. 2004; Wang et al. 2015). Since both liver and kidney are involved in MetS (Kaur 2014), we generated RNA-seq expression data from liver of LL rats and from kidney of all three strains and integrated these with relevant existing data including the level of regulatory conservation for three liver-specific transcription factors (CEBPA, FOXA1 and HNF4A, Stefflova et al. 2013) between rat and five related mouse species and strains. We show that the level of functional regulatory conservation can help select strain-specific haplotype blocks putatively associated with phenotypic differences among Lyon rats.

## Results

### 85% of strain-specific variation among Lyon rat strains is concentrated in less than 9% of the genome

To define haplotype blocks, we partitioned the rat genome into 10kb windows and calculated the number of strain-specific variants (SSVs) in each window relative to the reference rat genome assembly (see methods). We observed a bimodal distribution in the number of SSVs and used this distribution to define the resulting haplotype blocks as having either a high density of SSVs (High Variability Region, HVR) or a low density of SSVs (Low Variability Region, LVR) (see Methods, Figures 1 and S1).

**Figure 1.**
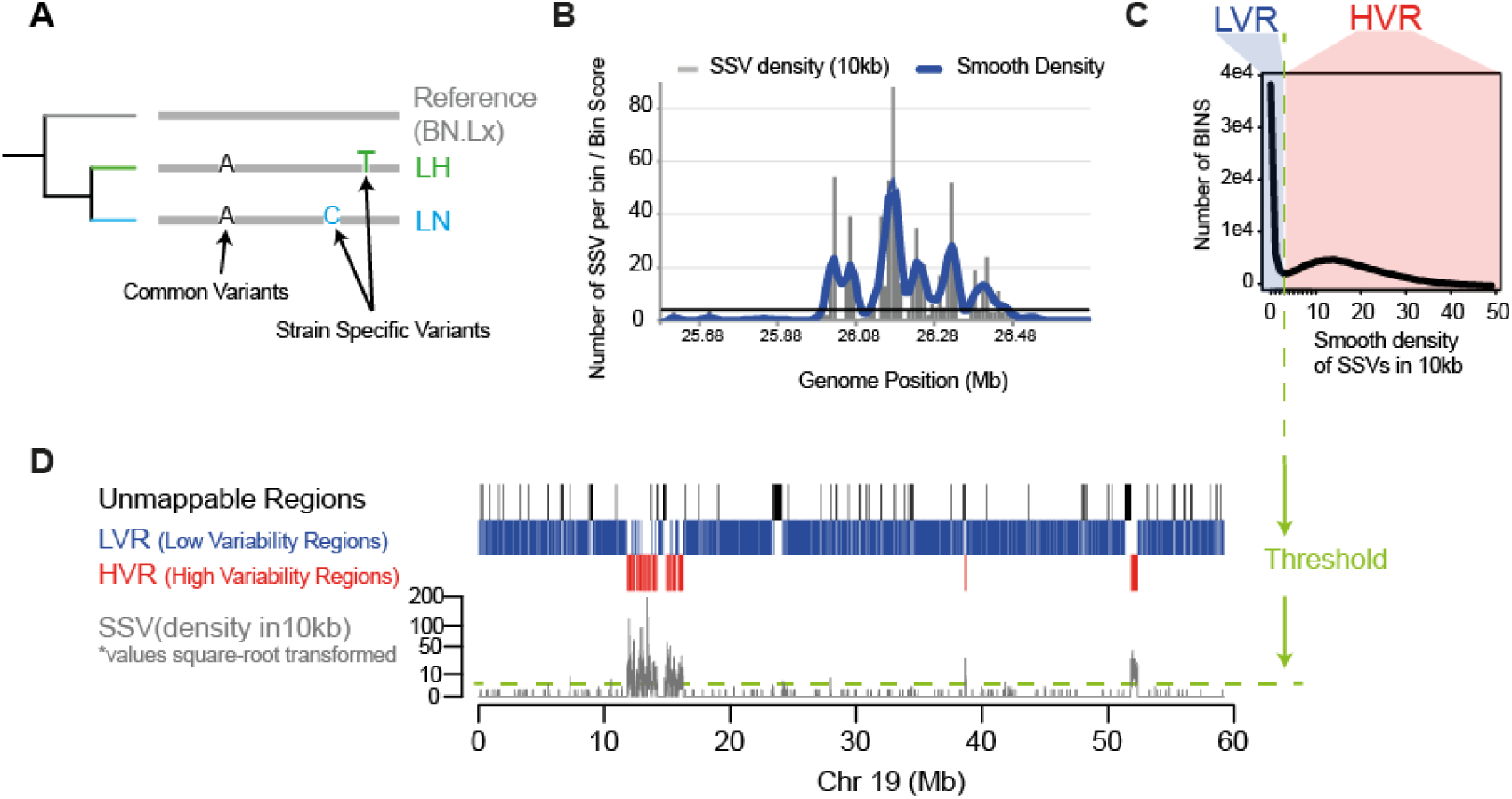
Procedure to identify genomic regions of interest based on the distribution of Strain-Specific Variants (SSV) across the genome. (A) SSVs were obtained for the pairwise comparisons between the two susceptible Lyon strains (LH or LL) relative to the control Lyon rat (LHvsLN and LLvsLN) and the reference rat genome (LHvsBN and LLvsBN). (B) Densities of SSVs were calculated in non-overlapping genome windows of 10kbp; we applied a smoothing algorithm to these densities (see Methods). (C) Distribution of the smoothed densities of SSVs obtained from LNvsBN and LLvsBN were used to calculate the threshold between HVRs and LVRs. Only those regions with at least three consecutive genome windows of the same type were considered. (D) HVR and LVR across chr19 for the hypertensive Lyon rat relative to the control Lyon rat (LHvsLN); regions showing poor mapping qualities (unmappable) were discarded from our analyses.

The distribution of SSVs across the genome was similar for the two pairwise comparisons of Lyon rats susceptible to MetS and obesity (LH and LL) and the Lyon Normotensive (LN) that is resistant (i.e. LHvsLN and LLvsLN, see Figures 2A, S3A, S4A). In both cases, the vast majority of strain-specific variants were concentrated in HVRs (LHvsLN: 84.96% and LLvsLN: 85.09%), and these regions only covered a small part of the genome (LHvsLN: 8.55% and LLvsLN: 7.10%) (Figures 2B, S3B, S4B and Table 1). These regions were partly overlapping: 42.6% of LHvsLN HVRs overlap with LLvsLN HVRs, while 51.2% of LLvsLN HVRs overlap with LHvsLN. SSV overlaps have similar fractions (Figure 2C). The fraction of the genome that we identify as highly strain-specific is similar to that obtained previously for these and other rat strains (see Atanur et al. 2013; Ma et al. 2014, Methods and Figure S3). HVRs characterise a substantial reduction in the portion of the genome that is most likely to be involved in MetS phenotypes and therefore form the primary focus of our subsequent analysis.

**Figure 2.**
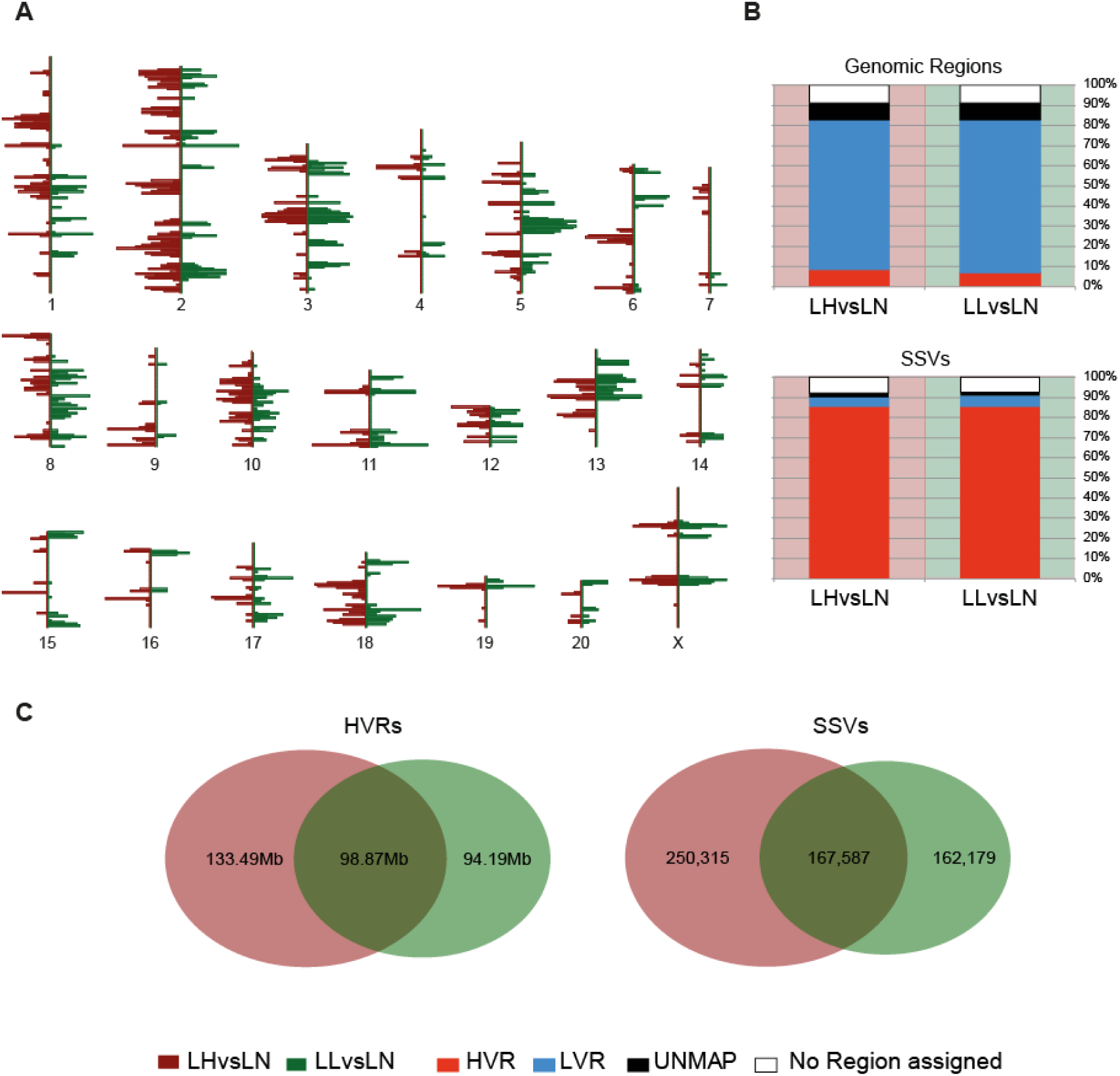
Genome distribution and characteristics of HVRs and SSVs. (A) Distribution and density across the whole genome of the HVRs obtained for LHvsLN (in brown) and LLvsLN (in green) rats. Figure modified from the Ensembl genome browser version 69. (B) Percentages of the rat genome and the strain-specific variants assigned to HVR, LVR and unmappable regions (UNMAP) for LHvsLN and LLvsLN. (C) Overlap of HVRs and SSVs for LHvsLN and LLvsLN comparisons.

**Table 1.**
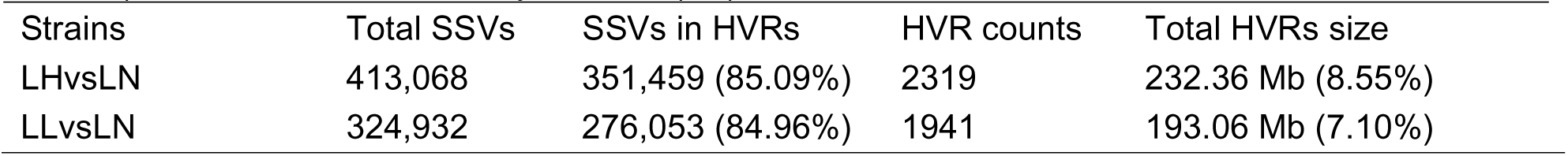
Descriptive statistics for the SSVs and HVRs obtained for the susceptible Lyon rats (LH and LL) relative to the resistant Lyon strain (LN).

| Strains | Total SSVs | SSVs in HVRs | HVR counts | Total HVRs size |
| --- | --- | --- | --- | --- |
| LHvsLN | 413,068 | 351,459 (85.09%) | 2319 | 232.36 Mb (8.55%) |
| LLvsLN | 324,932 | 276,053 (84.96%) | 1941 | 193.06 Mb (7.10%) |

### Evidence for the functionality of High Variability Regions in the Lyon rats associated with MetS

We then sought to determine if the HVRs preferentially contain features that could explain the phenotypic differences among Lyon rats by comparing them to other regions of the genome (see Methods and Figure 3). Specifically, we tested whether there is a significant enrichment in HVRs of the following elements: i) annotated genes, ii) genes associated with metabolic-related traits, iii) genes differently expressed among Lyon rats, iv) occupancy in rat of three liver-specific transcription factors, and v) regions orthologous to human variants associated by GWAS to obesity and metabolic traits.

**Figure 3.**
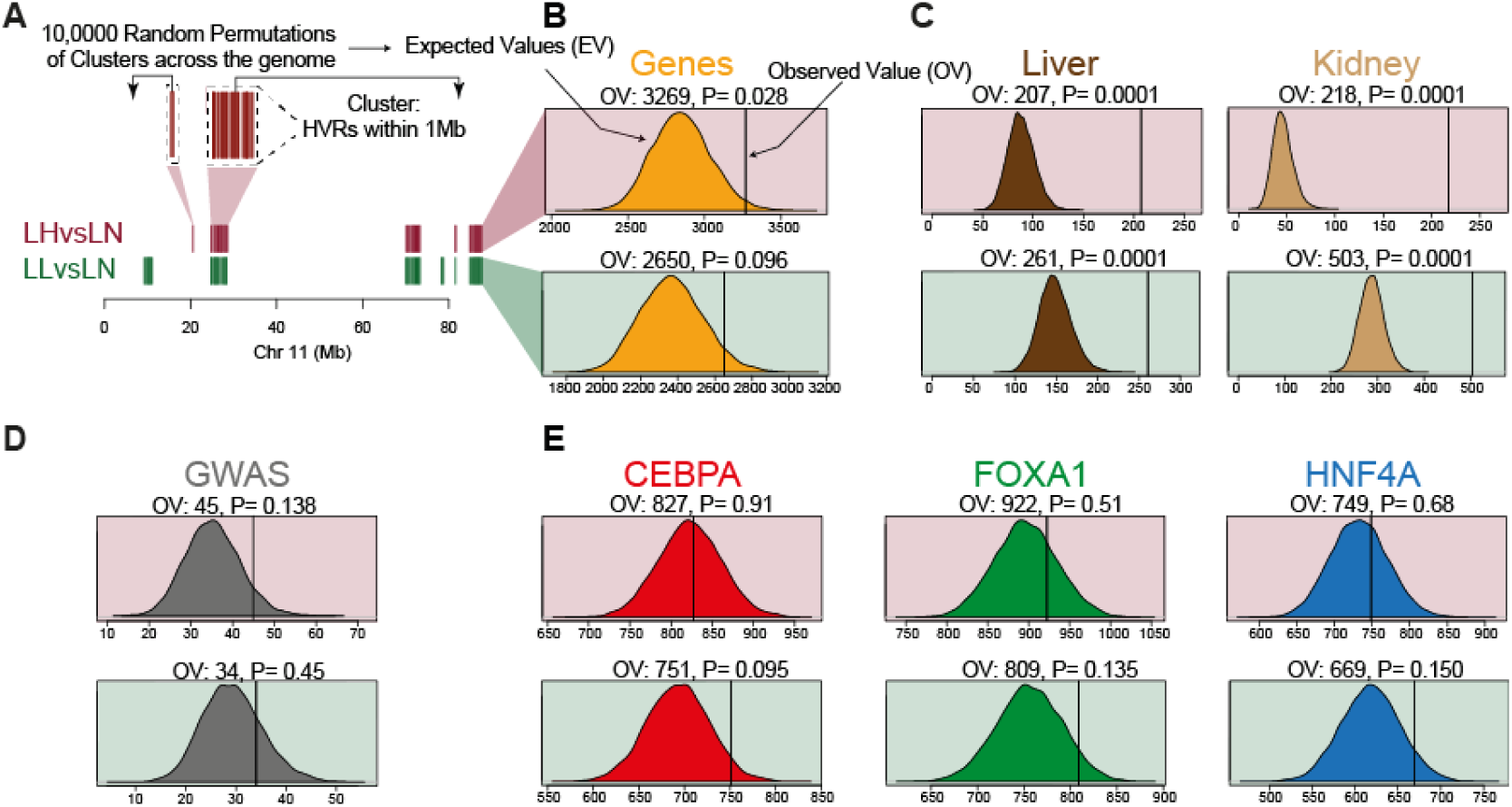
High Variability Regions as functional regions. (A) HVRs and Ensembl genes (orange) across chr11 for LHvsLN (red) and for LLvsLN (green). We tested for significant enrichment of HVRs with several genetic elements using permutation tests. We compared the observed values (OV) and the expected values (EV) to determine significance. Similar analyses was done for: (B) Ensembl annotated rat genes (v69) (C) Genes whose expression differed either in liver or in kidney between LH and LN strains, and between LL and LN. (D) Rat orthologues of human GWAS variants associated with metabolic traits. (E) Occupancy in BN rat strain of three liver-specific transcription factors (CEBPA, FOXA1 and HFN4A)

The number of Ensembl genes (Yates et al. 2016) that overlap at least one HVR was marginally greater than expected by chance and this overlap was significant for the LHvsLN comparison (p < 0.05), but not for LLvsLN (p > 0.05) (Figure 3B). Additionally, but only in the case of LLvsLN, there was a significant enrichment of genes associated with Type I diabetes mellitus (KEGG PATHWAY database, gene count: 25, p < 10^−9^, see results for ‘All HVRs’ in Figures S5 and S6 (DAVID web services v6.7, Huang da et al. 2009; Jiao et al. 2012). Gene enrichment in the HVRs was more significant when considering only the genes whose expression in either liver or kidney differed between LH and LN strains, and between LL and LN strains (see Methods and Table S1). (Figure 3C and Table S2).

We next considered whether the HVRs were enriched for either the occupancy of three specific transcription factors (HNF4A, CEBPA, and FOXA1) or the 418 rat orthologues of Human GWAS variants associated with obesity and metabolic traits. In both cases we did not observe a significant enrichment (Figures 3D-E, S7 and Table S3).

In summary, the observation that both annotated and differentially expressed genes are enriched in HVRs supports the hypothesis that HVRs harbour functional regions that could be responsible for phenotypic differences observed among the Lyon rats. However, given the overall genomic span of identified HVRs and the large number of SSVs both in coding and non-coding regions in the HVRs (see Table 1), these analyses on their own are inadequate to suggest either the causative genes or the causative variants influencing MetS or obesity across the whole genome.

### Liver-specific regulation data can prioritise regions of strain-specific variation in Lyon rats associated with MetS

We next integrated strain-specific genomic variation with available genomic regulatory data from tissue relevant to MetS in order to prioritise the HVRs using the occupancy and level of conservation of the three liver-specific transcription factors. We created subsets of HVRs characterised by occupancy of the factors and the level of conservation among mice and rats using a factor-specific Conservation Enrichment score (CE_f_, see Methods). Briefly, CE_f_ is the fraction of transcription factor binding events in a 10kb window that are conserved between rat and mouse for each transcription factor. The score was determined independently for each of the three factors (f = CEBPA, FOXA1, or HNF4A). Thus, for each transcription factor and for both the LHvsLN and LLvsLN comparisons, we created seven subsets of HVRs: ‘All HVRs’ including those without any binding event; ‘HVR w/TFBS’ with at least one TFBS regardless of conservation; and five subsets containing HVRs with CE_f_ greater than 0, 0.2, 0.4, 0.6 and 0.8, respectively. We then reassessed the evidence for functionally of these HVR subsets in a similar way to that done with the whole set of HVRs as above.

Enrichment of Ensembl rat genes in HVR subsets corresponded with the occupancy and level of conservation of the three liver-specific transcription factors (Figure 4A). With the exception of the subset of LLvsLN with all HVRs, all tests in the HVR subsets were statistically significant (p <0.05). For LHvsLN and for the three factors, maximum significance possible (p < 10^−3^) was obtained for the subset of HVRs with at least one TFBS (HVR w/TFBS), and for the subsets with CE_f_>0.0 and CE_f_>0.2. In the case of LLvsLN and for the three transcription factors, the maximum significance was obtained in the subset of HVRs with at least one TFBS, and HVR subset with CE_f_>0.0 (i.e. HVR subsets with the darkest colour in Figure 4A).

**Figure 4.**
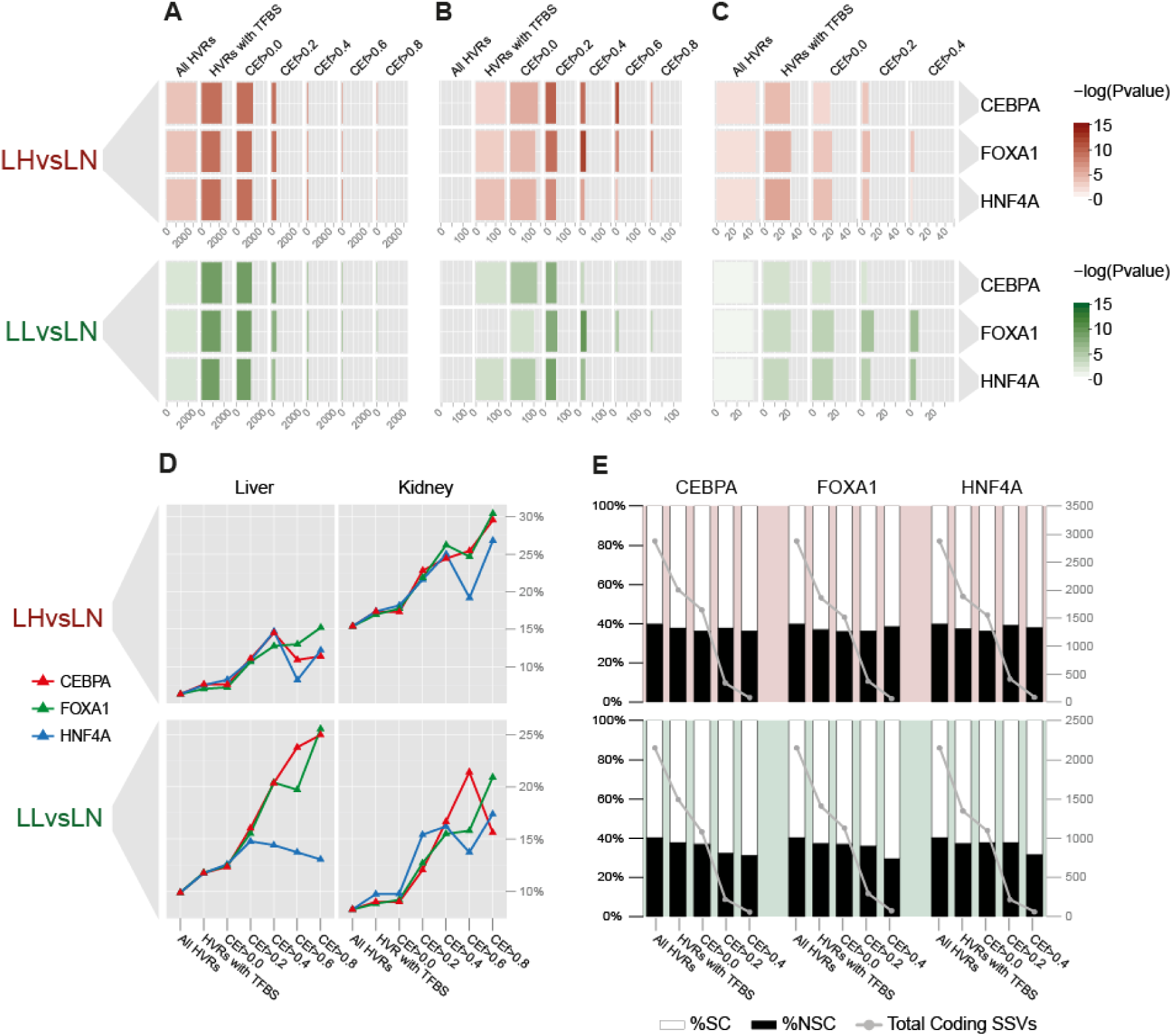
Conservation Enrichment (CE_f_) is correlated with functional enrichment. (A) Genes overlapping HVR subsets. The colour gradient shows the significance obtained from the enrichment analyses done in each of the HVR subsets. Bar sizes indicates the total number of genes overlapping each subset of HVRs. (B) Liver-genes defined by UP_TISSUE annotation. The functional annotation analyses were calculated by DAVID 6.7. Colour gradient and bar size defined as above. (C) Rat orthologues of human GWAS regions associated with obesity and metabolic traits. We excluded HVR subset with CE_f_>0.6 and CE_f_>0.8 due to no GWAS variants overlapping these HVR subsets. Colour gradient and bar size defined as above. (D) Ratio of genes differentially expressed in either liver or kidney, between LH and LN strains, and between LL and LN strains (y-axis) for each subset of HVRs. (E) Proportion of synonymous coding SSV (SC-SSV) and non-synonymous coding SSVs (NSC-SSV) in HVRs for each subset of HVRs.

As above, we analysed the functional annotation enrichment in the HVR subsets using DAVID (Huang da et al. 2009; Jiao et al. 2012) with KEGG PATHWAY (Kanehisa and Goto 2000; Kanehisa et al. 2014) and UP TISSUE (Uniprot Consortium 2015) databases (see Methods). In all cases (both LHvsLN and LLvsLN for the three liver-specific transcription factors) the term ‘liver’ from the UP TISSUE database, had the greatest accumulated significance across HVR subsets (see Figures S5 and S6). These results indicate that HVRs selected according to information from liver-specific regulation data are enriched in genes associated with liver function. Importantly, but as expected, this association was not evident without using genomic regulation data to select HVRs (see Figure 4B).

In the case of LHvsLN, the greatest enrichments in genes associated with liver function were obtained for the CE_CEBPA_>0.6 (p < 10^−5^), CE_FOXA1_>0.4 (p < 10^−5^) and CE_HNF4A_>0.2 (p < 10^−3^) subsets (HVR subsets with the darkest colour in Figure 4B). For LLvsLN, the greatest enrichments were obtained for CE_CEBPA_>0.2 (p < 10^−3^), CE_FOXA1_>0.4 (p < 10^−4^) and CE_HNF4A_>0.2 (p < 10^−3^) (Figure 4B).

For the analyses using KEGG_PATHWAY database, we did not find a consistent increase in significance associated with an increase in CE_f_, although we did identify some functional terms that were statistically significant (see Figures S5 and S6).

Genes differentially expressed between LH and LN strains in either liver or kidney (Table S1) are significantly enriched for all subsets of HVRs (Figure S9), regardless of which transcription factor is considered. The same is true for genes differentially expressed between the LL and LN strains. To compare the differences in enrichment between subsets, we computed the fraction of all Ensembl rat genes that are differently expressed for each HVR. In all cases, the fraction of differentially expressed genes was positively correlated with CE_f_ (Figure 4D). For example, for LHvsLN and data from liver, the fraction increased from 6% (‘all HVRs’ subset) to 15% (CE_FOXA1_>0.8), while for kidney, it increased from 15% (‘all HVRs’ subset,) to 30% (CE_CEBPA_>0.8; CE_FOXA1_>0.8). A similar pattern was observed for LLvsLN (Figure 4D).

We hypothesised that there may be a correlation between selection pressures leading to regulatory conservation as measured by CE_f_ and changes to the sequence of protein coding genes within the same sets of HVRs. The ratio of non-synonymous coding SSVs (NSC-SSVs) to synonymous coding SSVs (SC-SSVs) was therefore compared across HVR subsets (see Methods). Although we find relatively little difference in the ratio of the non-synonymous changes, especially for the case of the LHvsLN comparison, in the LLvsLN comparison, non-synonymous changes do appear to be depleted when HVRs have higher regulatory conservation (i.e. higher CE_f_) (Figure 4E). This may be the effect of simultaneous selection on both protein coding genes and regulatory networks for a subset of regions in the LL genome.

Finally, we looked for an enrichment of putative GWAS positive regions in HVR subsets by determining the orthologous location in rat of NHGRI-EBI GWAS Catalog SNPs associated with obesity and metabolic-related in humans (Welter et al. 2014) (specific terms listed in Table S3). The use of the regulatory information from liver-specific transcription factors identified significant enrichments of GWAS variants in relevant subsets of HVRs. For example, we found significant enrichments (i.e. p < 0.05) for LHvsLN in the subsets of HVRs w/ TFBS and CE_f_>0.2 for the three liver-specific factors, in CE_f_>0.0 for FOXA1 and HNF4A and in CE_f_>0.4 for FOXA1) (Figure 4C, Table S6). Regarding LLvsLN, we found significant enrichments for in HVRs w/TFBS, CE_f_>0.0, CE_f_>0.2, CE_f_>0.4 for both FOXA1 and HNF4A (Figure 4C, Table S6).

In summary, the use of CE_f_ (i.e. the conservation level between rat and mouse in the occupancy of three liver-specific transcription factors) is effective for selecting candidate regions involved in phenotypic differences between Lyon rats. In most cases, we observed an increase in statistical significance as a function of CE_f_. Moreover, the consideration of regulatory information was required to identify HVRs significantly enriched for genes associated with metabolic related-trait genes and enriched for human GWAS variants related to obesity and metabolic traits.

### Integrating results from the three liver-specific transcription factors to prioritise the strain-specific variation in Lyon rats associated with MetS

Given the observed stability of combinatorially bound transcription factors (Stefflova et al. 2013) and connection of these regions to human disease (Ballester et al. 2014), we assessed if the number of liver-specific transcription factors used to estimate the conservation level could more efficiently prioritise candidate HVRs. For this purpose, we used the HVR subsets with CE_f_>0 (i.e. all HVRs with at least one conserved TFBS).

We observed that conservation of more than one type of factor in a given HVR was common: 41% (LHvsLN) and 43% (LLvsLN) of the HVR CE_f_>0 regions had conserved peaks for all three of the liver-specific transcription factors (Figure 5A). We then partitioned the HVRs with conserved peaks by the diversity of factors that where conserved in the given HVR. Specifically, ‘HVR 1TF’ includes HVRs with one or more conserved TFBS from at least one factor; while ‘HVR 2TF’ and ‘HVR 3TF’ refer to HVRs with conserved TFBS from at least two or all three factors (i.e. HRV 3TF is a strict subset of HVR 2TF, which is strict subset of HVR 1TF). We then assessed these HVRs subsets to determine if an increased diversity of conserved liver-specific transcription factors is an effective method to prioritise HVRs.

**Figure 5.**
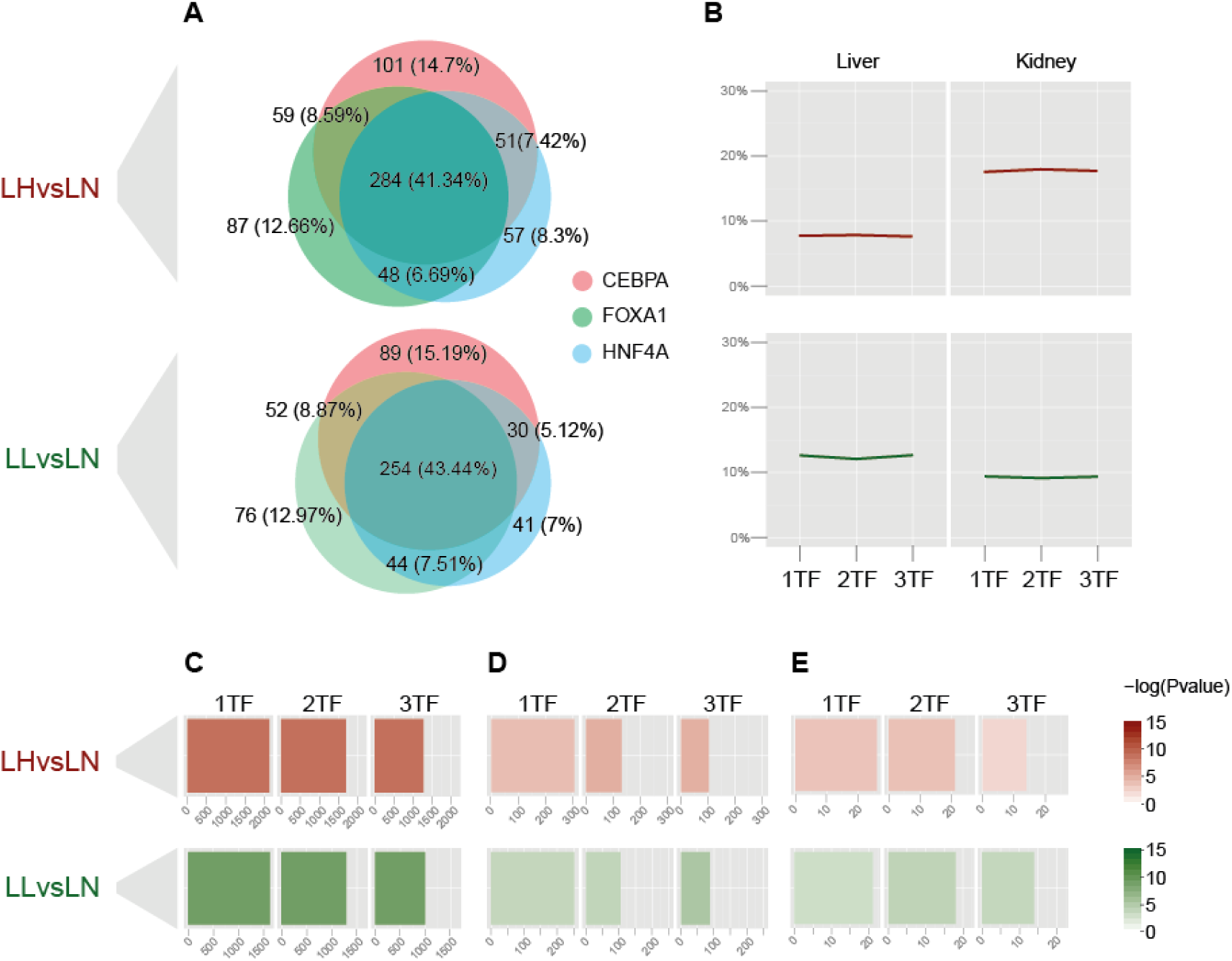
Integrating results from the three liver-specific transcription factors. (A) Venn diagrams of HVRs with at least one conserved binding site between rats and mice (HVR subsets with CE_f_>0) for the three liver-specific transcription factors. (B) Ratio of genes differentially expressed in either liver or kidney between LH and LN, and between LL and LN calculated for each subset of HVRs is not influenced by the number of bound factors. (C) Genes overlapping HVR subsets. The colour gradient shows the significance obtained from the enrichment analyses. Bar sizes indicates the total number of genes overlapping each subset of HVRs. (D) Liver-genes according to UP_TISSUE annotation. The functional annotation analyses were calculated by DAVID 6.7. Colour gradient and bar size defined as above. (E) Rat orthologues of human GWAS SNPs associated with obesity and metabolic traits. Colour gradient and bar size defined as above.

Although the presence of genes is significantly enriched in each of these HVR subsets, there are no differences among HVR 1TF, HVR 2TF and HVR 3TF: in all cases enrichment significances were equal to p = 10^−4^ (Figure 5C). Similar results were obtained when considering all liver-associated genes in HVRs (see methods) for LLvsLN (HVR TF1: gene count = 126, p < 0.05; HVR TF2: gene count = 105, p < 0.05; HVR TF3: gene count = 87, p < 10^−2^) and LHvsLN (HVR TF1: gene count = 153, p < 0.05; HVR TF2: gene count = 131, p < 10^−2^, HVR TF3: gene count = 102, p < 10^−2^) (Figure 5D).

HVRs with an increased diversity of conserved peaks were generally significantly enriched (permutation tests, p < 0.05) for orthologous regions of human GWAS SNPs except for the case of the HVR 3TF subset with the LHvsLN SSVs (Figure 5E and Table S7).

The significance of enrichments of genes differentially expressed in either liver or kidney was p < 10^−3^ for the subsets with at least one conserved peak for one, two and three liver-specific transcription factors, respectively, for both LHvsLN and LLvsLN strain comparisons. Considering the ratios of differentially expressed genes, we observed that they kept relatively constant across HVRs subsets as the number of liver-specific transcription factors with conserved peaks increased (Figure 5B).

These results suggest that knowledge of which TFBSs are conserved and whether a given region of the genome has conserved TFBSs from multiple factors may be effective in some situations at prioritising regions with strain specific variation involved with tissue specific functions. For example, we did not observe enrichments in HVRs associated with liver-expressed genes and the orthologous rat regions associated with human GWAS without using the conservation level as measured by CE_f_ (see above and Figure 4).

### Analysing the genes obtained from the selected High Variability Regions

To gain insight into genes from the prioritised HVRs that may be important for MetS or obesity, we focused on the most conserved and regulatorily complex HVR subset, i.e. the set containing at least one conserved peak for all three liver-specific factors (the HVR 3TF subset, see above).

We performed two analyses based on possible functional mechanisms underpinning the phenotypic differences among Lyon rats. First, we characterised those genes with non-synonymous coding strain-specific variants (NSC-SSVs; see Methods) overlapping the selected HVRs. Such variation would result in changes to the amino acid sequence that may be responsible for functional changes in the resulting proteins. Second, we identified those genes located near putative promoters in rat obtained from Villar et al. (2015) (see Methods) and with SSVs overlapping the selected HVRs. We assumed these SSVs might affect the expression of the proximal genes. For these analyses, we used only those genes expressed in liver as measured by RNA-seq data (FKPM > 1, see Methods).

We categorised the selected genes based on whether they were i) liver-specific genes (according to the UniProt tissue database, see Methods), ii) differentially expressed in liver and/or kidney when comparing the susceptible Lyon strains with the control Lyon strain, (see Methods); iii) overlapping human GWAS variants associated with obesity and metabolic traits overlapping the gene body in the case of genes with NSC-SSVs or overlapping the promoter in the cases of genes linked to promoters with SSVs (see Table 2 and Supplementary Material).

**Table 2.** Number of rat genes and human orthologues expressed in liver of the susceptible Lyon rats (LH or LL) and associated with coding or non-coding (promoters) strain-specific variation overlapping the selected subsets of HVRs obtained from the two strain comparisons (i.e. HVR with at least one conserved peaks for three liver-specific transcription factors). Numbers of human orthologues with evidenced associations with insulin resistance, dyslipidaemias and obesity, are shown between parentheses.

|  | LHvsLN |  | LLvsLN |  |
| --- | --- | --- | --- | --- |
|  | Rat genes | Human orthologues | Rat genes | Human orthologues |
| **Coding Variation** |  |  |  |  |
| *Genes with NSC-SSVs* | 133 | 111 (1,1,14) | 96 | 78 (0,0,8) |
| *Genes with GWAS<sup>‡</sup> and NSC-SSVs* | 3 | 2 (0,0,2) | 3 | 3 (0,0,2) |
| *Liver-genes<sup>§</sup> with NSC-SSVs* | 15 | 9 (0,1,4) | 10 | 6 (0,0,1) |
| *Dif-liver<sup>\*</sup> genes with NSC-SSVs* | 14 | 12 (0,0,0) | 17 | 16 (0,0,1) |
| *Dif-kidney<sup>\*</sup> genes with NSC-SSVs* | 32 | 25 (0,0,4) | 9 | 7 (0,0,0) |
| **Promoters** |  |  |  |  |
| *Genes assigned to promoters with SSVs* | 232 | 206 (7,3,32) | 189 | 164 (4,4,18) |
| *Genes assigned to promoters with GWAS<sup>‡</sup> and SSVs* | 1 | 1 (0,0,1) | 1 | 1 (0,0,1) |
| *Liver-genes<sup>§</sup> assigned to promoters with SSVs* | 35 | 30 (2,1,9) | 26 | 26 (1,1,4) |
| *Dif-liver<sup>\*</sup> assigned to promoters with SSVs* | 38 | 33 (1,1,8) | 26 | 26 (1,1,3) |
| *Dif-kidney<sup>\*</sup> assigned to promoters with SSVs* | 52 | 44 (0,1,7) | 13 | 13 (0,1,1) |

<sup>‡</sup> Human GWAS variants associated with Obesity and Metabolic traits in the NHGRI-EBI GWAS catalogue
<sup>§</sup> Genes expressed specifically in liver according to UniProt tissue database and accessed by using DAVID web services.
<sup>\*</sup> Genes differently expressed in liver or in kidney when comparing the susceptible Lyon rats (LH or LL) with the control Lyon rat (LH).

We also analysed the genes associated by published evidence to three symptoms showed by the LH strain (insulin resistance, dyslipidaemias) and by the LH and LL strains (obesity) plus two symptoms not obviously present in these strains as control (heart disease and Alzheimers), using DisGeNET (v4.0, Piñero et al. 2015; Piñero et al. 2016) (see Methods). For this analysis, we used the corresponding human orthologues of the selected rat genes because the DisGeNET data is mainly for human (see Methods). A total of 7,542 and 7,520 rat genes had human orthologues and were expressed in liver in the LH and LL strains, respectively. DisGeNET identifies a small number of these genes as associated with the metabolic syndrome phenotypes and, as expected, these gene sets are highly similar for the LH and LL strains with approximately 140 (1.9%), 110 (1.5%) and 800 (10.6%) genes associated with insulin resistance, dyslipidaemia and obesity, respectively in each strain.

There were 173 protein-coding genes expressed in liver of LH or LL strains with at least one NSC-SSV in the selected HVRs. Of these 173 genes, 144 had identified human orthologues and can thus be compared with the DisGeNET data. This set of genes was not significantly enriched for published associations to obesity, insulin resistance or dyslipidaemias, (see Table 2, Supplementary Material).

A larger number of genes were one-to-one associated with putative active promoters including 3,865 and 3,864 genes that were expressed in livers of LH and LL strain rats and had human orthologues, respectively. Of the 3,865 genes with active promoters from the LH strain, a total of 85 (2.2%), 66 (1.7%) and 425 (11%) were associated with insulin resistance, dyslipidaemias and obesity, respectively. The number for LL are similar: 86 (2.2%), 65 (1.7%), 417 (10.8%) for the associations to insulin resistance, dyslipidaemias and obesity, respectively. Only a fraction of these promoters had SSVs in the selected HVRs: 206/3865 (5.3%) for LH and 164/3864 (4.2%) for LL. Thirty-two of the genes assigned to promoters with SSVs overlapping the selected HVRs in the LH strain were associated with obesity (15.5%, Fisher’s exact test: p < 0.05). There were no significant enrichments for insulin resistance or dyslipidaemias in the LH strain or any significant associations in the LL comparison (see Table 2, Table S10 and supplementary Material).

We found no significant enrichments for the two symptoms used as control in either the comparison to genes with at least NSC-SSV in the selected HVRs (Fisher’s exact tests: all p-values > 0.08) or to genes one-to-one assigned to promoters with SSVs overlapping the selected HVRs (Fisher’s exact tests: all p-values > 0.05).

Of the set of 32 genes responsible for the significant enrichment for obesity in the LHvsLN comparison (Figure S10), the gene with most published evidence of association with obesity was the insulin receptor gene *Insr* (ENSRNOG00000029986) (Table S9); *Cat* (ENSG00000121691) was the human gene of that list assigned to the promoter with the greatest number of SSVs (58 SSVs) overlapping the HVRs.

### Discussion

In this study, we have used the level of functional regulatory conservation between related species to prioritise genomic regions whose patterns of genome variation suggest that they are involved in phenotypic differences in a model of obesity and metabolic syndrome, the Lyon rat strains.

As a first step, we characterised haplotype blocks by density of strain-specific variants for the two comparisons between the susceptible Lyon strains with respect to the resistant Lyon strain (i.e. LHvsLN and LLvsLN). In agreement with similar analyses (Atanur et al. 2013; Ma et al. 2014), most of these variants were concentrated in a small part of the genomes, which we termed High Variability Regions (HVRs). Next, we classified the HVRs according to conserved occupancy between rat and mice for three liver-specific transcription factors. Functional enrichment of selected HVRs was evident for those genetic elements where a significant enrichment was found in the whole HVR sets. Importantly, our approach revealed associations between HVRs with liver-genes and with rat orthologues of human GWAS linked to obesity and metabolic traits.

We also searched genes associated with genomic variation linked to two selected sets of HVRs, one from each strain comparison; namely, those sets with haplotype blocks having at least one conserved peak among rat and mice for each of the three liver-specific transcription factors (i.e. ‘HVR 3TF’ subset). In these two subsets, we determined those genes with non-synonymous strain-specific variants and genes assigned to promoters with strain-specific variation overlapping the selected haplotype blocks. We reported a list of these selected genes where we included additional information coming from functional analyses and supporting the association of these genes with human GWAS for obesity and metabolic traits and with traits in the susceptible Lyon strains (insulin resistance, dyslipidaemias and obesity) (Supplementary Material). We found a significant enrichment of liver-expressed genes associated with obesity that were assigned to promoters with strain-specific variation overlapping the selected haplotype block obtained from LHvsLN.

Ma, et al characterised the blocks with a high density of variants that are unique in the Lyon strains in order to fine-map Quantitative Trait Loci (QTL) for MetS previously identified in these rat strains (Ma et al. 2014). As result, the candidate QTL regions were narrowed by 78%. By focusing their analyses to coding variants in the QTL on rat chromosome 17, they reduced the number of candidate genes to 27. We found that 3 of these genes had non-synonymous strain-specific variation overlapping the most stringent HVR 3TF subset (see Table S10), however none of these genes were assigned to promoters with strain-specific variation overlapping HVR 3TF regions. More recently, Wang, et al, reported 17 candidate genes involved in the phenotypic differences between LH and LN Lyon rats (Wang et al. 2015). We found that only one of Wang et al.’s genes held non-synonymous variation overlapping an HVR 3TF region (see Table S11). In addition, none of the genes identified by Wang, et al, were assigned to promoters with strain-specific variation overlapping the HVR 3TF regions (Table S10).

The gene *RGD1562963* (ENSRNOG00000039379), which encodes a protein similar to chromosome 6 open reading frame 52 (C6ORF52), is the only one reported by both studies in the previous paragraph. It is suggested to be the most likely eQTL driver gene involved in phenotypic differences between LH and LN strains (Wang et al. 2015). *RGD1562963* is cis-regulated by an eQTL hotspot on chromosome 17 and is predicted to affect 100 of 278 trans-eQTL genes (Wang et al. 2015). While this gene was not linked to strain-specific variation overlapping the strict HVR 3TF subset, *RGD1562963* is associated with the less restrictive HVRs 2TF subset obtained from LHvsLN comparison. Moreover, *RGD1562963* was the only gene from the Ma et al. and Wang et al. lists with both non-synonymous coding and promoter assigned SSVs (Figure 6). This result would suggest that *RGD1562963* has been under positive selection during the phenotype-driven derivation of this strain and gives support to the predicted role of *RGD1562963* affecting susceptibility in LH rats for the Metabolic syndrome reported by Ma et al. and Wang et al. The identification of *RGD1562963* by our complementary method further supports its role in the phenotype and lends additional validation to our general approach.

**Figure 6.**
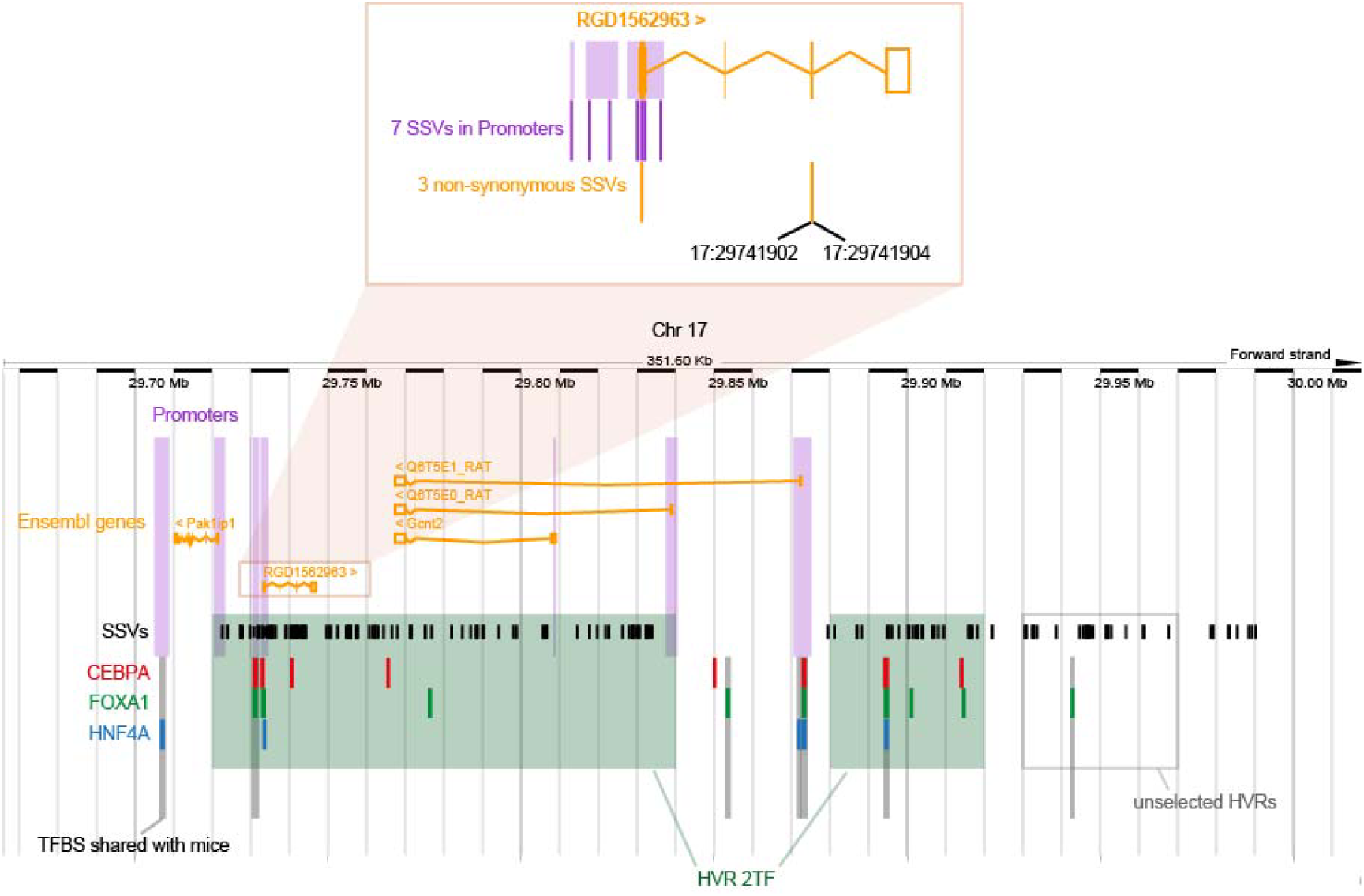
The genomic region of *RGD1562963* (*ENSRNOG00000039379*) Non-synonymous coding SSVs and promoter SSVs linked to *RGD1562963* overlapping HVRs from the LHvsLN strain comparison are highlighted. Promoter splitting in the inset image was due to conversion from the RGSC3.4 coordinate system. Image modified from Ensembl genome browser (*Rattus norvegicus* version 69.34 (RGSC3.4) Chromosome 17: 29,665,942 - 30,017,536).

Our results demonstrate both the potential and the limitations of using the level of functional regulatory conservation to prioritise genomic regions potentially associated with phenotypic differences among Lyon rats. This approach would be most easily extended to other systems with similar breeding histories including other rat strains and mice strains. Importantly, it is not needed to generate data from many individuals like QTL and eQTL approaches and allows the use information already available, as conservation in regulatory elements between rat and mice.

## Methods

### Determination of High Variability Regions, Low Variability Regions and Unmappable Regions

#### Genomic sequences and Single Nucleotide Variants

We used existing whole genome alignments (*ENA accession: ERP002160*) and single-nucleotide variants (available from the Rat Genome Database) of the three Lyon strains (LH, LL and LN) that were generated by Atanur et al. (2013) in comparison to the BN reference genome (RGSC-3.4, Gibbs et al. 2004).

##### Strain-Specific Variant (SSV)

We called a SSV for a given strain as a genomic position with an allele that is not present in the strain used as reference (Figures 1A and S1A). Firstly, we obtained SSVs for Lyon strains compared to the BN reference genome RGSC-3.4 and the resulting sets of SSVs are referred to as LHvsBN, LLvsBN and LNvsBN for the SSVs specific to the LH, LL and LN strains, respectively. These comparisons were used to calculate the threshold for the different types of genomic regions (see below and Figures 1C and S1C). Secondly, we obtained SSVs for the two pairwise comparisons of Lyon rats that are susceptible to MetS and obesity phenotypes relative to the strain that is resistant (LHvsLN and LLvsLN). In these cases, we called a SSV as a genomic position with at least one allele that is not present in both LN and the reference BN genome. By doing this, we discard from LH and LL genomes the genetic variation shared with LN strain, which we assume are not associated with MetS (Figures 2, S3 and S4). Furthermore, in a similar way as done for LN strain by comparing it to LH and LL strains (LNvsLH, LNvsLL); these comparisons were used as controls of our approach, because we expected to not find any association between LN strain SSVs and MetS (see Figures S3 and S4).

In a previous study with the Lyon rats, Ma et al. (2014) considered a SSV as any position that differed between the two strains that were being compared regardless of whether the position was variable with respect to the reference BN genome (indicated as LH+LN, LN+LH in Figure S3 and LL+LN and LN+LL in Figure S4). In our case, we considered a SSV only if the allele both differed from the BN reference genome and was also in the strain that was used as query and not present in the strain used as control. For example, in Figure S1, our approach considers only the G in the LNvsLN comparison, while Ma, et al, would have included both the G and the C. This different criterion allowed us to remove from our LHvsLN and LLvsLN analyses those genomic regions specific to LN, and which are likely not associated with the phenotypic differences associated with MetS among Lyon rats (see Figures S3, S4). Other differences from methodology used by Ma, et al, were i) we did not discard the roughly 5% of SSVs that were called heterozygous by Atanur et al. (2013); and ii) we did not use those genome regions with a low estimated accessibility (see below).

##### Smooth Density of SSVs

For downstream analyses, we used a weighted sliding window approach—triangular smoothing—to calculate the number of SSVs in non-overlapping 10kb genome windows (Figures 1B and S1B). This method smoothes differences among windows that were caused by the genome compartmentalization. For a given window in the genome at position *x*, we calculated the smoothed density of SSVs as the following floating mean with weights:

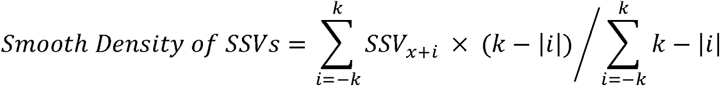

where *SSV_x+i_* is the number of Strain Specific Variants in the window with position *x+i*, and *k* is the number of neighbouring windows up and downstream used for smoothing. We use k=3 in our analyses empirically (data not shown) because this value gives a clear distinction between two types of genomic regions (see below and Figures 1C and S1C).

#### Genomic regions

##### High Variability Regions and Low Variability Regions

The smoothed density of SSVs in genome windows between two rat strains shows a bimodal distribution (Figures 1C and S1C). The left peak in the bimodal distribution contains regions of the genome identical by descent, with low a density of SSVs (Low Variability Region, LVR). The right peak contains regions of the genome that are divergent between the two strains with a high density of SSVs. A distinct valley separates the two peaks, which we used as a threshold to differentiate HVR and LVR. We calculated this threshold for the three comparisons between the Lyon strain rats and the reference rat genome (RGSC-3.4). In all three cases the threshold obtained was three (Figure S1C); that is, windows with a smoothed SSV density greater than three variants in 10kb were classified as HVR, and windows whose smoothed SSV density was less than or equal to three were classified as LVR. Only those regions with at least three consecutive genome windows of the same type were considered for further analyses (Figure 1D).

##### Unmappable regions

We performed two analyses on the BAM files to estimate the parameters to characterise the non-accessible genome regions of the Lyon strains. Firstly, we obtained the distribution of mapping qualities (i.e. -10 log_10_ Pr(mapping position is wrong), http://samtools.github.io/hts-specs/) by using QualityScoreDistribution.jar from Picard tools (v1.81(1299), http://picard.sourceforge.net) with the option VALIDATION_STRINGENCY=LENIENT (Figure S2A). Secondly, we calculate genome coverage per base by using *genomeCoverageBed* form *Bedtools* (v2.17.0, Quinlan and Hall 2010) with default parameters (Figure S2B). According to results obtained from the later analysis, we considered a region as unmappable when at least three consecutive windows with an average mapping quality less then or equal to 30 and/or with an average coverage greater then or equal to 100 (Figures 1D, 2, S3B and S4B).

### Animals

LH/MRrrcAek, LN/MRrrcAek, and LL/MRrrcAek rats were bred and maintained in an approved animal facility at the University of Iowa on a 12-hour light-dark cycle and provided food and water *ad libitum*. Male offspring were used in this study. All animal protocols were approved by the Institutional Animal Care and Use Committee (IACUC) at the University of Iowa. The rats were phenotyped and tissues collected as previously described (Wang et al. 2015). Briefly, at three weeks of age the rats were weaned onto normal chow (Teklad 7913 - Harlan Teklad NIH-31 irradiated, 18% protein, 6% fat). At 15 weeks of age they were switched to a 4% NaCl diet (Teklad 7913 modified with 4% NaCl) until they were humanely euthanized with CO_2_ at 18 weeks of age after an overnight fast. Tissues were collected and stored in RNAlater (Life Technologies, Grand Island, N.Y.) at -80°C for subsequent RNA extraction.

### Gene expression

#### RNA-seq data

RNA was isolated from liver and kidney tissue using standard TRIzol methods (Chomczynski and Sacchi 1987). RNA quality was measured (BioAnalyzer 2100, Agilent Technologies, Santa Clara, CA, USA), using an RIN threshold of 7. Libraries were prepared using TruSeq RNA Sample Preparation Kits v2 (Illumina, San Diego, CA) according to manufacturer’s instructions. RNA sequencing was performed on an Illumina HiSeq 2000, with paired-end, 50 bp cycles, at the Iowa Institute of Human Genetics - Genomics Division. Six samples were multiplexed per lane, yielding approximately 30 million reads per sample. All data consisted of six biological replicates for LH and LL liver and five biological replicates for LL liver and LH, LL, and LN kidney. Sequence data from LH and LN liver was previously described (Wang et al. (2015); GSE50027). Remaining sequence data created for this study has been deposited in the ArrayExpress database at EMBL-EBI (www.ebi.ac.uk/arrayexpress) under accession number E-MTAB-5939.

We analysed the read quality using FASTQC software (v 0.10.1 http://www.bioinformatics.babraham.ac.uk/projects/fastqc/). Reads were trimmed by using Trimmomatic (v0.32, Bolger et al. 2014) if the Phred score of any base was below 25 (LEADING:25 TRAILING:25). We used reads with at least 36 bases (MINLEN:36) and only those paired reads that remained after trimming.

#### Gene expression analyses

We estimated differential gene expression between LH and LN, and between LL and LN. TopHat (v2.0.13, Trapnell et al. 2009) was used to map reads to the rat reference genome (RGSC-3.4). Read alignments with more than two mismatches were discarded (*--read-mismatches 2*). We also used the option *--no-novel-juncs* to look for reads across junctions already annotated. The Cufflinks package (v2.2.1, Trapnell et al. 2010) was used to assemble transcripts separately for each replicate. We used *cuffmerge* to merge the transcript assemblies from replicates to be analysed, and finally we used *cuffdiff* to find differently expressed genes and with Benjamini-Hochberg corrected False Discovery Rates (FDR) of 0.05. Thus, we obtained the differently expressed genes either in liver or in kidney between LH and LN, and between LL and LN (see Table S1). We also used the FPKM (Fragments per kilobase of exon per million of fragments mapped) values obtained from these analyses to get the list of genes that were expressed in the livers of the LH and LL strains. We considered a gene expressed in liver if its FPKM was greater than 1.0.

### Regulation data: liver-specific transcription factors

We used the liver ChIP-seq datasets generated by Stefflova et al. (2013) for BN rat strain (ArrayExpress accession: E-MTAB-1414) and for five mouse species/strains (*Mus musculus* (strains C57BL/6J and AJ), *Mus caroli* and *Mus castaneus* and *Mus spretus,* ArrayExpress accession: E-MTAB-1414). The dataset comprised two biological replicates for each species/strain and for three liver-specific transcription factors (CEBPA, HNF4A and FOXA1). Reads were aligned using BWA (Li and Durbin 2009) with default parameters. Peak locations were called by SWEMBL (https://github.com/stevenwilder/SWEMBL). Final peak sets contained peaks present in both biological replicates.

#### Conservation of occupancy of three liver-specific transcription factors between rat and mouse strains

We compared peaks generated from ChIP-seq datasets among the five mouse species and the rat. We used only the genomic regions present in the BLAST-Z alignment between mouse and rat available in Ensembl (v59, Yates et al. 2016) and using the NCBI37 mouse genome as references for comparing datasets from the different species. We considered as conserved peaks between rat and mouse, the overlapping peaks between rat and at least one mouse species/strain. Coordinates of conserved peaks were converted to the rat genome reference (RGSC-3.4). For each HVR and liver-transcription factor, we calculated its Conservation Enrichment score (CE_f_) as the number of conserved peaks in 10kb for a given transcription factor.

### Permutation tests

Permutation tests were used to find significant enrichments in HVRs. We clustered HVRs within an empirically defined distance of 1Mb because HVRs have a non-uniform distribution across the genome (see Figure 3A) and we assume nearby HVRs are regulatorily non-independent. Clusters were randomly permutated across the whole genome by using the command *shuffle* from BEDTools (v2.22.0, Quinlan and Hall) and accessed from *pybedtools* (v0.6.9, Dale et al. 2011); the relative coordinates of HVRs inside of clusters were maintained. We estimated the distribution of expected values by calculating either the total number or average of genetic elements overlapping the set of HVRs inside of the shuffled cluster for each permutation. We performed 10,000 permutations in each test. Significance of the enrichment of the genetic element in HVRs was obtained by calculating the two-tailed p-value according to this formula:

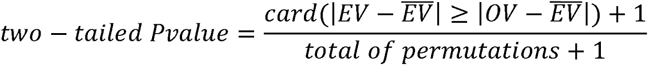

where *OV* is the value obtained from the observed HVRs, and *EV* is the expected value calculated from each of the 10,000 sets of permuted HVRs (see Figure 3A). The minimal p-value possible with 10,000 permutations is 1x10^−4^.

### Functional analyses of HVRs

#### Ensembl genes overlapping HVRs

We used the set of Ensembl genes from Ensembl (v69, Yates et al. 2016) for the BN reference genome RGSC-3.4. We used a permutation test (see above) to determine if genes overlapped HVRs more often than expected by chance. We calculated the number of genes overlapping at least one HVR in the observed permutated sets (Figure 3B).

#### Differentially expressed genes overlapping HVRs

We tested if genes that were differentially expressed between LH and LN and between LL and LN overlapped HVRs more often than expected by chance (Figure 3C). We used permutation test for these analyses (see above). We used the list of genes differentially expressed that were obtained from RNA-seq data (Table S1). We calculated the number of these genes that overlapped at least one HVR in the observed and permutated sets of HVRs (Figure 3C and Table S2).

#### Gene-annotation enrichment analysis of HVRs

We analysed if there was a functional enrichment associated with metabolic or obesity phenotypes for the genes overlapping at least one HVR that were obtained from LHvsLN (‘All HVRs’ in Figure S5) and LLvsLN (‘All HVRs’ in Figure S6). We tested for this enrichment by using DAVID web services v6.7 (python client, Huang da et al. 2009; Jiao et al. 2012) for KEGG PATHWAY (Kanehisa and Goto 2000; Kanehisa et al. 2014) and UP TISSUE (Uniprot Consortium 2015) databases (release/download date: Sep 2009, https://david.ncifcrf.gov/content.jsp?file=update.html). We used DAVID v6.7 (Sep 2009) for our analyses rather than DAVID v6.8 (October 2016), because most of the data used in our study (gene annotation, SNPs and occupancies of liver-transcription factors) are based on the RGSCv3.4 assembly, which is also that used by DAVID v6.7. DAVID v6.8 uses the Rnor 6.0 assembly and differences in the gene sets and/or gene nomenclature between these two rat assemblies create inconsistencies that affect the accuracy of our results (data not shown). In addition, although the KEGG PATHWAY resource was updated in DAVID 6.8, the UP TISSUE dataset, which we used to report the expected association between the term liver and the level of functional regulatory conservation (see Results section), was not updated in DAVID v6.8 (in both versions UP TISSUE is dated Sep 2009). We recognise that in their recent paper, Wadi et al. (2016) showed that the use of out-dated gene annotation prevents the identification of all significant terms in enrichment analyses. However, in our case, even when using DAVID v6.7, we found significant results and the expected correlation between gene enrichment and the level of functional regulatory conservation. Thus, the DAVID supporting database that we use are largely the same between v6.7 and v6.8, it is more important for us to be consistent on the assembly and gene set for our analysis.

#### Liver-specific transcription factor overlapping HVRs

We also tested if the number of peaks in rat overlapping HVRs (Figures 3E and S7) and the average CE_f_ (Figure S8) observed for each one of the three liver-specific transcription factors were significantly greater that that expected by chance. We used permutation tests (see above) for these analyses. For the observed values, we used either the total number of peaks overlapping the HVRs or the average CE_f_ for a given transcription factor. For the expected values, we calculated the two latter values for each one of the 10,000 permuted sets of HVRs (see above).

#### Human GWAS variants associated with metabolic traits overlapping HVRs

We obtained from the NHGRI-EBI GWAS catalogue (Welter et al. 2014) the list of SNPs associated with obesity and metabolic-related traits in humans (search terms used in Table S3). SNPs coordinates were converted from the GRCh38 human assembly to the rat RGSC-3.4 assembly using mapping from GRCh38 to Rno6.0 and then from Rno6.0 to Rno5.0 and RGSC-3.4. All conversions used the Ensembl Perl API and the Ensembl assembly converter software (v87, Yates et al. 2016). As with other genetic elements analysed, we then used permutation tests to determine if there was a significant enrichment of rat orthologous positions for these GWAS variants overlapping HVRs. For the observed value, we used the total number of GWAS variants overlapping HVRs. The expected value was calculated as the number of GWAS variants overlapping each one of the 10,000 sets of permuted sets of HVRs.

### Selection of HVRs according to CE_f_

For downstream analyses, we created seven subsets of HVRs according to the occupancy for the three liver-specific transcription factors and their CE_f_ for each on each one of the three liver-specific transcription factors (CECEBPA, CEFOXA1 and CEHNF4A): all HVRs; HVRs with at least one peak (HVR w/TFBS); HVRs with CE_f_ greater than 0 (i.e. HVRs with at least one conserved peak), and HVRs with CE_f_ greater than 0.2, 0.4, 0.6, 0.8 respectively (Table S4 and Table S5 show sizes and number of SSVs of HVR subsets). We analysed each one of the subsets of HVRs in a similar way to that used for the full set of HVRs as described above. Then, we compared the results obtained in each analysis across the subsets of HVRs (Figure 4). Specifically, we analysed the enrichment in HVRs for i) Ensembl genes, ii) gene annotation from DAVID (UP TISSUE and KEGG PATHWAY databases), iii) differentially expressed genes in liver and kidney (Table S1) and iv) rat orthologues of human GWAS variants associated with obesity and metabolic-related traits (Tables S3·and S6). Additionally, we also tested if the proportion of non-synonymous coding SSVs (NSC-SSVs) and synonymous coding SSVs (SC-SSVs) in HVRs differed between the subsets of HVRs. For this, we estimated the effect of SSVs in HVRs by using the Ensembl Variant Effect Predictor (VEP) tool (standalone perl script v2.7 associated with Ensembl v69, McLaren et al. 2016). We considered in the analyses those NSC-SSVs whose most severe effect was ‘*missense_variant*’, ‘*stop_gained’* or ‘*stop_lost*’.

### Selection of HVRs by the number of liver-specific transcription factors with conserved peaks

Three subsets of HVRs were created according to how many liver-specific transcription factors had conserved peaks: the ‘*HVR 1TF*’ subset included HVRs with conserved peaks for at least one liver-specific factor, the ‘*HVR 2TF*’ subset had HVRs with conserved peaks for at least two factors, and the ‘*HVR 3TF*’ subset had HVRs with conserved peaks for all three liver-specific factors (Figure 5A and Table S8). We compared the functionality among these HVRs subsets to test the importance of the number of transcription factors used to define the conservation level (Figure 5). We analysed each one of these three subsets of HVRs in a similar way as used for the full set of HVRs and for the HVRs subsets created with different conservation levels as described in the previous section. Specifically, we compared the enrichment in HVRs among ‘*HVR 1TF*’, ‘*HVR 2TF*’ and ‘*HVR 3TF*’ subsets for i) Ensembl genes, ii) gene annotation from DAVID (liver term of UP_TISSUE database), iii) differentially expressed genes in liver and kidney and iv) rat orthologues of human GWAS variants associated with obesity and metabolic-related traits.

### Analyses of SSVs of the selected subsets of HVRs

For these analyses, we selected the subset of HVRs that had at least one conserved peak between rat and mouse strains/species for all three of the liver-specific transcription factors (i.e. ‘*HVR 3TF’* subset) as they show enrichment for most of the functional elements and because of the observed stability of combinatorially bound transcription factors (Stefflova et al. 2013). We limited our analysis to the genes that were both expressed in liver of LH or LL (FKPM > 1) and associated with coding or non-coding strain-specific variation.

#### Non-synonymous coding SSVs (NSC-SSVs) in the selected subsets of HVRs

We assessed the effect of the SSVs on the protein by using the VEP tool (standalone perl script v2.7, McLaren et al. 2016). We considered in the analyses those SSVs classified as non-synonymous variants and whose most severe effect was ‘*missense_variant*’, ‘*stop_gained’* or ‘*stop_lost*’.

#### SSVs of the selected subsets of HVRs sited in promoters

Positions of putative promoters in Rat were obtained from Villar et al. (2015). These authors characterised promoters and enhancers by using modifications to histone 3 lysine 27 (H3K27ac) and histone 3 lysine 4 (H3K4me3). Active promoters are marked by H3K4me3 and H3K27ac, while active enhancers are regions marked by H3K27ac (Villar et al. 2015). Coordinates were converted from the Rnor5.0 assembly to the RGSC-3.4 assembly using the Ensembl assembly converter software (v80, Yates et al. 2016). Genes were assigned to promoters if the gene’s transcription start site (TSS) overlapped or was within 5kb downstream of the promoter. Only one-to-one gene-promoter assignations were used for our analysis.

### Association between metabolic diseases and genes

Genes associated with the three metabolic-related symptoms showed by LH and LL strains (i.e. insulin resistance, dyslipidaemias and obesity) was obtained from DisGeNET (v4.0, Piñero et al. 2015; Piñero et al. 2016). DisGeNET is a platform integrating information on associations between genes and human diseases from public data sources and literature. We analysed those genes expressed in liver and with either the selected NSC-SSV or assigned to selected promoters with SSVs. DisGeNET analysis used the human orthologous genes of the selected rat genes with homology determined by the Ensembl Perl API (v69, Yates et al. 2016). Only the human orthologous genes with rat homology annotated as *‘one2one’* or *‘apparently one2one’* were used. From DisGeNET, we searched for disease gene associations using relevant Unified Medical Language System Concept Unique Identifiers (UMLS^®^ CUIs, insulin resistance: C0021655, dyslipidaemias: C0242339 and obesity: C0028754). We also included two additional diseases not shown by the susceptible Lyon strains as controls for our analyses (heart diseases: C0018799 and Alzheimers: C0002395).

For each of the five diseases, we compared, using Fisher’s exact test, the counts of rat genes expressed in liver with NSC-SSVs overlapping the selected subsets of HVRs and human orthologues associated with that disease with the total number of rat genes expressed in liver and human orthologues associated with the disease. A similar comparison was done for genes assigned to promoters with SSVs overlapping the selected HVRs. In this case, the total number of human orthologues of rat genes expressed in liver was limited to those that were one-to-one assigned to promoters.

## Acknowledgments

We thank John Marioni for helpful discussions. RNA-seq data were obtained at the Genomics Division of the Iowa Institute of Human Genetics, which is supported, in part, by the University of Iowa Carver College of Medicine. The research leading to these results has received funding from the European Community’s Seventh Framework Programme (FP7/2007-2013) under grant agreement Nº HEALTH-F4-2010-241504 (EURATRANS), and from NIH R01 HL089895 (AEK) and R21 DK089417 (AEK). We acknowledge additional support from the Wellcome Trust (WT108749/Z/15/Z) and the European Molecular Biology Laboratory.

## Authors contributions

The study was designed and supervised by DT and PF. DMG did all analysis except for the definition of HVRs and LVRs, which was done by DDdS. RNA-seq data was generated by AEK and MCJM. DMG and PF wrote the paper with input from all authors.

## Related Figure

**Figure S1.-** Procedure to identify genomic regions of interest based on the distribution of SSV across the genome.

**Figure S3.-** HVR/LVR approach in LHvsLN.

**Figure S4.-** HVR/LVR approach in LLvsLN.

**Figure S5.-** Gene-annotation enrichment analyses performed with DAVID v6.7 in HVRs obtained for the comparison LHvsLN.

**Figure S6.-** Gene-annotation enrichment analyses performed with DAVID v6.7 in HVRs obtained for the comparison LLvsLN.

**Figure S7.-** HVRs and occupancy of the occupancy for three liver-specific transcription factors.

**Figure S5.-** Gene-annotation enrichment analyses performed with DAVID v6.7 in HVRs obtained for the comparison LHvsLN.

**Figure S6.-** Gene-annotation enrichment analyses performed with DAVID v6.7 in HVRs obtained for the comparison LLvsLN.

**Figure S8.-** HVRs and conservation between rat and mice for the occupancy of three liver-specific transcription factors.

**Figure S9.-** Gene expression and level of conservation between rat and mice for the occupancy of the three liver- specific transcription factors.

## References

Adams DJ, Doran AG, Lilue J, Keane TM. 2015. The Mouse Genomes Project: a repository of inbred laboratory mouse strain genomes. Mamm Genome 26: 403-412.

Aitman T, Dhillon P, Geurts AM. 2016. A RATional choice for translational research? The Company of Biologists Ltd.

Aitman T, Petretto E, Behmoaras J. 2010. Genetic Mapping and Positional Cloning. In Rat Genomics, Vol 597 (ed. Anegon I), pp. 13-32. Humana Press.

Atanur SS, Diaz AG, Maratou K, Sarkis A, Rotival M, Game L, Tschannen MR, Kaisaki PJ, Otto GW, Ma MC et al. 2013. Genome sequencing reveals loci under artificial selection that underlie disease phenotypes in the laboratory rat. Cell 154: 691-703.

Ballester B, Medina-Rivera A, Schmidt D, Gonzalez-Porta M, Carlucci M, Chen XT, Chessman K, Faure AJ, Funnell APW, Goncalves A et al. 2014. Multi-species, multi-transcription factor binding highlights conserved control of tissue-specific biological pathways. Elife 3.

Baud A, Hermsen R, Guryev V, Stridh P, Graham D, McBride MW, Foroud T, Calderari S, Diez M, Ockinger J et al. 2013. Combined sequence-based and genetic mapping analysis of complex traits in outbred rats. Nat Genet 45: 767-775.

Berman BP, Nibu Y, Pfeiffer BD, Tomancak P, Celniker SE, Levine M, Rubin GM, Eisen MB. 2002. Exploiting transcription factor binding site clustering to identify cis-regulatory modules involved in pattern formation in the Drosophila genome. Proc Natl Acad Sci U S A 99: 757-762.

Bilusic M, Bataillard A, Tschannen MR, Gao L, Barreto NE, Vincent M, Wang T, Jacob HJ, Sassard J, Kwitek AE. 2004. Mapping the genetic determinants of hypertension, metabolic diseases, and related phenotypes in the lyon hypertensive rat. Hypertension 44: 695-701.

Bolger AM, Lohse M, Usadel B. 2014. Trimmomatic: a flexible trimmer for Illumina sequence data. Bioinformatics 30: 2114-2120.

Cheng Y, Ma Z, Kim B-H, Wu W, Cayting P, Boyle AP, Sundaram V, Xing X, Dogan N, Li J. 2014. Principles of regulatory information conservation between mouse and human. Nature 515: 371-375.

Chomczynski P, Sacchi N. 1987. Single-step method of RNA isolation by acid guanidinium thiocyanate-phenol-chloroform extraction. Anal Biochem 162: 156-159.

Consortium GP. 2015. A global reference for human genetic variation. Nature 526: 68-74.

Cuppen E. 2005. Haplotype-based genetics in mice and rats. Trends Genet 21: 318-322.

Dale RK, Pedersen BS, Quinlan AR. 2011. Pybedtools: a flexible Python library for manipulating genomic datasets and annotations. Bioinformatics 27: 3423-3424.

Dupont J, Dupont JC, Froment A, Milon H, Vincent M. 1973. Selection of three strains of rats with spontaneously different levels of blood pressure. Biomedicine 19: 36-41.

Dwinell MR, Lazar J, Geurts AM. 2011. The emerging role for rat models in gene discovery. Mamm Genome 22: 466-475.

Encode Project Consortium, Bernstein BE, Birney E, Dunham I, Green ED, Gunter C, Snyder M. 2012. An integrated encyclopedia of DNA elements in the human genome. Nature 489: 57-74.

Gibbs RA Weinstock GM Metzker ML Muzny DM Sodergren EJ Scherer S Scott G Steffen D Worley KC Burch PE et al. 2004. Genome sequence of the Brown Norway rat yields insights into mammalian evolution. Nature 428: 493-521.

Hermsen R, de Ligt J, Spee W, Blokzijl F, Schafer S, Adami E, Boymans S, Flink S, van Boxtel R, van der Weide RH et al. 2015. Genomic landscape of rat strain and substrain variation. BMC Genomics 16: 357.

Huang da W, Sherman BT, Lempicki RA. 2009. Bioinformatics enrichment tools: paths toward the comprehensive functional analysis of large gene lists. Nucleic Acids Res 37: 1-13.

Jacob HJ. 2010. The rat: a model used in biomedical research. Methods Mol Biol 597: 1-11.

Jiao X, Sherman BT, Huang DW, Stephens R, Baseler MW, Lane HC, Lempicki RA. 2012. DAVID-WS: a stateful web service to facilitate gene/protein list analysis. Bioinformatics 28: 1805-1806.

Kanehisa M, Goto S. 2000. KEGG: kyoto encyclopedia of genes and genomes. Nucleic Acids Res 28: 27-30.

Kanehisa M, Goto S, Sato Y, Kawashima M, Furumichi M, Tanabe M. 2014. Data, information, knowledge and principle: back to metabolism in KEGG. Nucleic Acids Res 42: D199-205.

Kaur J. 2014. A comprehensive review on metabolic syndrome. Cardiol Res Pract 2014: 943162.

Kimura M. 1968. Evolutionary rate at the molecular level. Nature 217: 624-626.

King JL, Jukes TH. 1969. Non-Darwinian evolution. Science 164: 788-798.

Li H, Durbin R. 2009. Fast and accurate short read alignment with Burrows-Wheeler transform. Bioinformatics 25: 1754-1760.

Lindsey JR, Baker HJ. 2006. Chapter 1 - Historical Foundations. In The Laboratory Rat (Second Edition), doi:http://dx.doi.org/10.1016/B978-012074903-4/50004-2 (ed. Franklin MASHWL), pp. 1-52. Academic Press, Burlington.

Lowdon RF, Jang HS, Wang T. 2016. Evolution of Epigenetic Regulation in Vertebrate Genomes. Trends Genet 32: 269-283.

Lowe WL, Reddy TE. 2015. Genomic approaches for understanding the genetics of complex disease. Genome Res 25: 1432-1441.

Ma MCJ, Atanur S, Aitman T, Kwitek A. 2014. Genomic structure of nucleotide diversity among Lyon rat models of metabolic syndrome. BMC Genomics 15: 197.

Mack KL, Nachman MW. 2016. Gene Regulation and Speciation. Trends Genet.

Mashimo T, Serikawa T. 2009. Rat resources in biomedical research. Curr Pharm Biotechnol 10: 214-220.

McLaren W, Gil L, Hunt SE, Riat HS, Ritchie GR, Thormann A, Flicek P, Cunningham F. 2016. The Ensembl Variant Effect Predictor. Genome biology 17: 1.

Moreno-Moral A, Petretto E. 2016. From integrative genomics to systems genetics in the rat to link genotypes to phenotypes. Disease Models & Mechanisms 9: 1097-1110.

Nica AC, Dermitzakis ET. 2013. Expression quantitative trait loci: present and future. Phil Trans R Soc B 368: 20120362.

Pai AA, Gilad Y. 2014. Comparative studies of gene regulatory mechanisms. Curr Opin Genet Dev 29: 68-74.

Pennacchio LA, Rubin EM. 2001. Genomic strategies to identify mammalian regulatory sequences. Nat Rev Genet 2: 100-109.

Piñero J, Bravo À, Queralt-Rosinach N, Gutiérrez-Sacristán A, Deu-Pons J, Centeno E, García-García J, Sanz F, Furlong LI. 2016. DisGeNET: a comprehensive platform integrating information on human disease-associated genes and variants. Nucleic Acids Res: gkw943.

Piñero J, Queralt-Rosinach N, Bravo À, Deu-Pons J, Bauer-Mehren A, Baron M, Sanz F, Furlong LI. 2015. DisGeNET: a discovery platform for the dynamical exploration of human diseases and their genes. Database 2015.

Quinlan AR, Hall IM. 2010. BEDTools: a flexible suite of utilities for comparing genomic features. Bioinformatics (Oxford, England) 26: 841-842.

Romero IG, Ruvinsky I, Gilad Y. 2012. Comparative studies of gene expression and the evolution of gene regulation. Nat Rev Genet 13: 505-516.

Saar K, Beck A, Bihoreau MT, Birney E, Brocklebank D, Chen Y, Cuppen E, Demonchy S, Dopazo J, Flicek P et al. 2008. SNP and haplotype mapping for genetic analysis in the rat. Nat Genet 40: 560-566.

Sassolas A, Vincent M, Benzoni D, Sassard J. 1981. Plasma lipids in genetically hypertensive rats of the Lyon strain. J Cardiovasc Pharmacol 3: 1008-1014.

Shibata Y, Sheffield NC, Fedrigo O, Babbitt CC, Wortham M, Tewari AK, London D, Song L, Lee B-K, Iyer VR. 2012. Extensive evolutionary changes in regulatory element activity during human origins are associated with altered gene expression and positive selection. PLoS Genet 8: e1002789.

Shimoyama M, De Pons J, Hayman GT, Laulederkind SJ, Liu W, Nigam R, Petri V, Smith JR, Tutaj M, Wang S-J. 2014. The Rat Genome Database 2015: genomic, phenotypic and environmental variations and disease. Nucleic Acids Res: gku1026.

Stefflova K, Thybert D, Wilson MD, Streeter I, Aleksic J, Karagianni P, Brazma A, Adams DJ, Talianidis I, Marioni JC et al. 2013. Cooperativity and rapid evolution of cobound transcription factors in closely related mammals. Cell 154: 530-540.

Trapnell C, Pachter L, Salzberg SL. 2009. TopHat: discovering splice junctions with RNA-Seq. Bioinformatics 25: 1105-1111.

Trapnell C, Williams BA, Pertea G, Mortazavi A, Kwan G, van Baren MJ, Salzberg SL, Wold BJ, Pachter L. 2010. Transcript assembly and quantification by RNA-Seq reveals unannotated transcripts and isoform switching during cell differentiation. Nat Biotechnol 28: 511-515.

UniProt Consortium. 2015. UniProt: a hub for protein information. Nucleic Acids Res 43: D204-212.

Villar D, Berthelot C, Aldridge S, Rayner TF, Lukk M, Pignatelli M, Park TJ, Deaville R, Erichsen JT, Jasinska AJ et al. 2015. Enhancer evolution across 20 mammalian species. Cell 160: 554-566.

Villar D, Flicek P, Odom DT. 2014. Evolution of transcription factor binding in metazoans - mechanisms and functional implications. Nat Rev Genet 15: 221-233.

Vincent M, Boussairi EH, Cartier R, Lo M, Sassolas A, Cerutti C, Barres C, Gustin MP, Cuisinaud G, Samani NJ et al. 1993. High blood pressure and metabolic disorders are associated in the Lyon hypertensive rat. J Hypertens 11: 1179-1185.

Voigt B. 2010. Rat strain repositories. Methods Mol Biol 597: 323-331.

Wadi L, Meyer M, Weiser J, Stein LD, Reimand J. 2016. Impact of outdated gene annotations on pathway enrichment analysis. Nat Methods 13: 705-706.

Wang J, Ma MCJ, Mennie AK, Pettus JM, Xu Y, Lin L, Traxler MG, Jakoubek J, Atanur SS, Aitman TJ et al. 2015. Systems biology with high-throughput sequencing reveals genetic mechanisms underlying the metabolic syndrome in the Lyon hypertensive rat. Circ Cardiovasc Genet 8: 316-326.

Ward LD, Kellis M. 2012. Interpreting noncoding genetic variation in complex traits and human disease. Nat Biotechnol 30: 1095-1106.

Welter D, MacArthur J, Morales J, Burdett T, Hall P, Junkins H, Klemm A, Flicek P, Manolio T, Hindorff L et al. 2014. The NHGRI GWAS Catalog, a curated resource of SNP-trait associations. Nucleic Acids Res 42: D1001-1006.

Wittkopp PJ, Kalay G. 2012. Cis-regulatory elements: molecular mechanisms and evolutionary processes underlying divergence. Nat Rev Genet 13: 59-69.

Wong ES, Thybert D, Schmitt BM, Stefflova K, Odom DT, Flicek P. 2015. Decoupling of evolutionary changes in transcription factor binding and gene expression in mammals. Genome Res 25: 167-178.

Yates A, Akanni W, Amode MR, Barrell D, Billis K, Carvalho-Silva D, Cummins C, Clapham P, Fitzgerald S, Gil L. 2016. Ensembl 2016. Nucleic Acids Res 44: D710-D716.

Yau AC, Holmdahl R. 2016. Rheumatoid arthritis: identifying and characterising polymorphisms using rat models. Disease Models & Mechanisms 9: 1111-1123.

